# NT-mini, a recombinant tool for the study of Neurotrypsin functionality

**DOI:** 10.1101/2020.10.01.322396

**Authors:** Anselmo Canciani, Cristina Capitanio, Serena Stanga, Silvia Faravelli, Pascal Kienlen-Campard, Federico Forneris

## Abstract

Neurotrypsin (NT) is a highly specific nervous system multi-domain serine-protease best known for its selective processing of the potent synaptic organiser agrin. Its enzymatic activity is thought to influence processes of synaptic plasticity, with its deregulation causing accelerated neuromuscular junction (NMJ) degeneration or contributing to forms of mental retardation. Something which, based on the available literature, likely stems from NT-based regulation of agrin signalling. However, dissecting the exact biological implications of NT-agrin interplay is difficult, owing to the scarce molecular detail regarding NT activity and NT-agrin interactions. The difficult recombinant production of NT in its catalytically competent form is at the base of these limitations, and is currently constraining a more detailed molecular, biochemical and structural characterisation of the NT-agrin system. We have developed a novel strategy to reliably produce and purify a truncated but catalytically competent variant of NT called NT-mini. The characterisation of our construct highlighted almost wild type-like behaviour with comparable specificity, and conservation of modulation by calcium and heparin despite the lack of several accessory domains. With the data obtained from NT-mini it was then possible to identify NT’s heparin-binding domain, and discover a novel putative Zinc-based modulation of NT. Additionally, NT-mini allowed us to investigate the effect of NT activity on myotube formation in controlled cell-based experiments, evidencing a negative impact on myoblast fusion dependant on enzymatic activity. Collectively, this shows the viability of NT-mini as a model to study NT functionality, allowing to expand both *in-vitro* and “*in-cellulo*” investigations and providing a foundation to unravel the molecular underpinnings and biological significance of NT-agrin interactions.

## Introduction

Synapses are responsible for the vital processes of signal transmission mediating both neuro-muscular communication and higher brain functions. The correct organization of processes underlying synaptic formation and maintenance is critical, and relies on coordinated signalling pathways integrating an array of mechanical, molecular, electrical and chemical signals^1–3^. In this context, proteolytic processing of synaptic organizers by specific proteases can serve as a potent tool to spatio-temporally modulate those pathways. This can also function as a mechanism to ensure synchronized communication and organization of pre- and postsynaptic architectures^2,4^. Several proteases have been described as contributing to such regulatory mechanisms, with differing degrees of specificity^5^. Among those Neurotrypsin^6^ (NT), also known as prss12, motopsin^7^ and leydin^8^, stands out for its specificity^9^, and its possible biological roles^10,11^ that would make it a common modulator of central (CNS) and peripheral nervous system (PNS) synaptic plasticity.

Discovered in the late 90’s, this nervous system multi-domain cysteine-rich serine-protease was initially described on the basis of predictive sequence analyses^6,7,12^. Produced predominantly by neurons of the CNS and PNS, NT expression peaks in the perinatal period, in a time-frame often associated with intense synaptogenesis and synaptic rearrangement. In adult organisms NT expression is maintained at a lower level, and remains highest within the CNS^1,13^. Interestingly, NT was found to accumulate in synaptic vesicles and to be released in the extracellular environment in an activity-dependant manner^14^. When secreted in the synapse NT cleaves its only known substrate, the large proteoglycan agrin, at two distinct sites (α and β), resulting in the generation of two specific fragments of 22 and 90 kDa. This process has been associated to synaptic rearrangement events influencing CNS synaptic plasticity^10,14^, and to the destabilisation of the nerve muscle contacts known as neuromuscular junctions (NMJs)^11,15^. Coordination of pre- and postsynaptic signalling seems to regulate NT activity at the CNS, hinting at a possible role in processes of activity driven synaptic plasticity and long-term potentiation that underlie learning and memory^4,10,16^. In line with NT’s contributions to synapse formation, deregulation of its activity was seen to affect CNS plasticity, and likely contribute to NMJ related neurodegenerative disorders. NT inactivation leads to a form of non-syndromic mental retardation, while its overactivity induces an accelerated aging and a destabilization of NMJs associated to sarcopoenia (a muscle wasting disease of the elderly)^11,17,18^. Interestingly, NT might even play a role in other major CNS neurodegenerative disorders, such as Alzheimer’s disease (AD), as its expression and activity were tentatively linked to Presenilins^17^ that are key players in AD onset and progression.

NT’s domain composition is inferred from the original sequence analyses, which highlighted the presence of a proline-rich N-terminal region followed by; 1 kringle (Kr) domain, 3-4 scavenger receptor cysteine-rich (SRCR1-4) domains, a predicted Furin-recognised zymogen activation site, and a final (C-terminal) catalytic serine-protease (SP) domain^6,12^. Experimental data mostly describes the catalytic activity of NT, providing a broad outline of its properties. NT activity was reported to be strongly calcium-dependant, most active in pseudo-physiological conditions, and subject to modulation by glycosaminoglycans such as heparin^19,20^. Furthermore, NT displays a stunningly high specificity, to the point that it is reported to only have one substrate: the large proteoglycan agrin^5,21^. While data regarding NT’s catalytic activity has allowed to infer not only the role but also several properties of its SP domain, the function of the remaining domains is still matter of debate. Of its 4-5 accessory domains (Kr-SRCR3/4), only the Kr and murine SRCR3 domains have been characterised and associated to putative functions^22,23^. The former was associated to the binding of a seizure related protein of the CNS (sez-6)^24^, while the latter was found to directly mediate binding to calcium^23^ essential to NT’s activity^19^.

Despite the relative amount of information supporting NT’s possible biological roles, specific knowledge regarding its molecular architecture/function is scarce. This is an intrinsic limitation to the detailed understanding of NT functionality, and particularly in how its interplay with agrin underpins its role in physiological and pathological processes. Thus, producing recombinant NT would be a strong asset to gain insight into NT’s function. However, the recombinant production of this multi-domain cysteine-rich protein is notoriously difficult, requiring complex techniques (murine hybridomas) yielding modest sample amounts^19^. A methodology which is hardly compatible with the avid requirements of protein structure studies, or those associated to extensive biochemical characterisation.

We attempted to address these limitations by developing a novel strategy for the recombinant production and purification of a catalytically competent fragment of human NT, as well as its inactive Ser825Ala mutant. Characterisation of these recombinant proteins, with biochemical and biophysical methods, allowed us to observe a behaviour analogous to that of the full-length NT, and expand upon some of the previously reported features regulating NT’s activity. This work offers new insights into NT-agrin interplay, but also provides a set of appealing tools to further investigate NT functionality in cell-based experiments.

## Results

### Production and purification of NT-mini/NT*-mini

In line with previously published data^19^, obtaining an enzymatically active human NT construct was particularly challenging. The full-length protein proved too intractable for production, thus requiring the engineering of an extensive library of shorter variants derived from the original cDNA. Of these, only a single construct, termed NT-mini (Gly497-Leu875), bearing the catalytically active SP domain, was successfully produced in a HEK293 mammalian cell suspension system. Purification of NT-mini from the cell culture media was obtained via a “mixed-mode” affinity chromatography approach based on immobilised metal ion (IMAC) and heparin affinity. In order to avoid the partial zyomgen-enzyme activation reported for the full murine NT^19^, the activation process was “enhanced upstream” by co-transfecting NT-mini with the pro-protein convertase Furin^25^. The final result was a highly pure and homogeneous sample almost entirely present in the active two-chain enzyme conformation (Fig. 1B), with an estimated yield of 1 milligram per litre of cell suspension culture.

**Figure 1 -.**
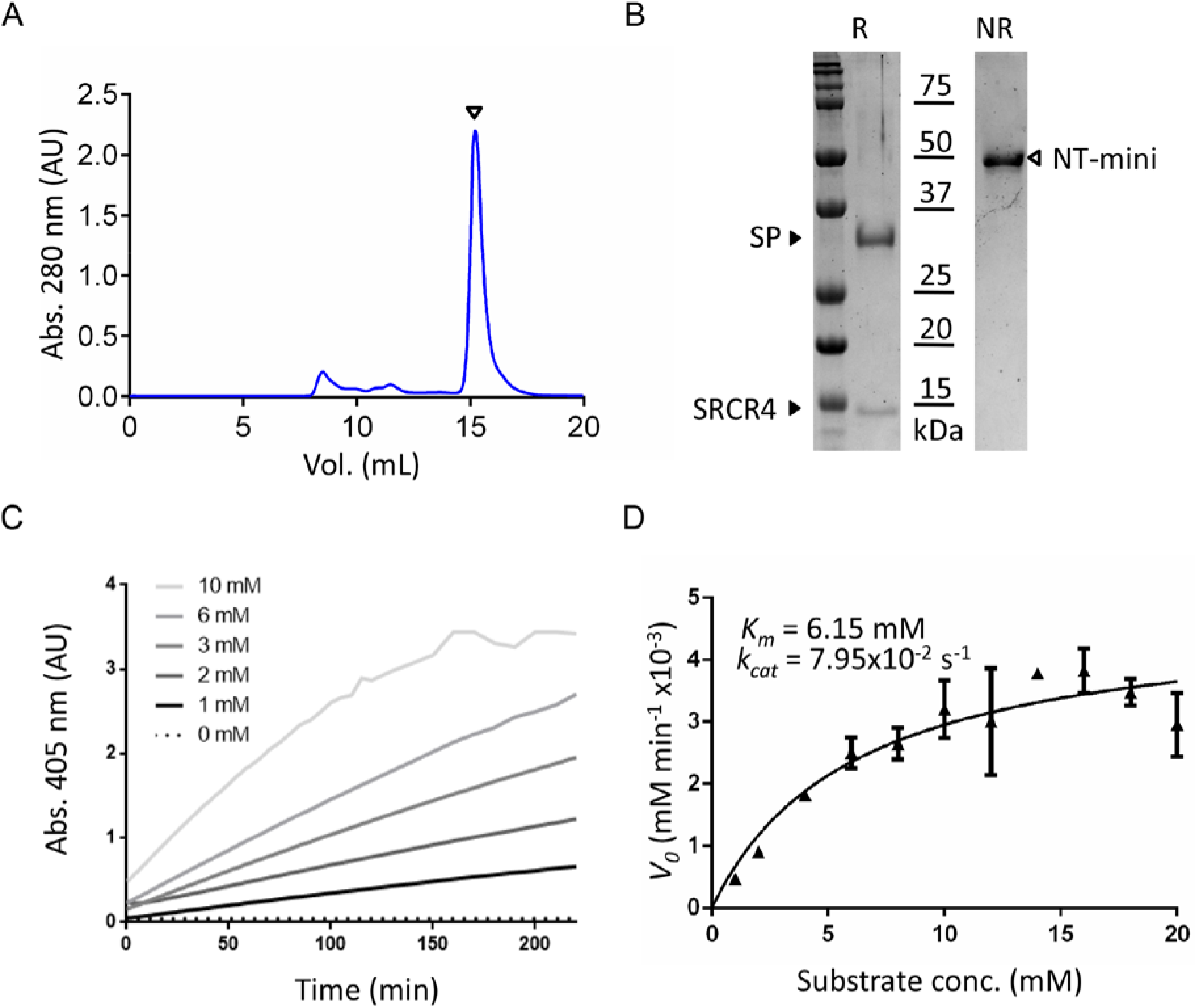
Summary of NT-mini purification and characterization with synthetic peptides. (A) Size exclusion chromatography (SEC) representative of the final purification step for NT-mini. The protein is visible as a single sharp peak (white triangle) well separated from other higher MW contaminants. (B) SDS-PAGE gel representative of the final purified product. The two-chain active form of NT-mini (white arrow) is visible as a single band close to the 50 kDa MW marker in non reducing conditions (NR). In reducing (R) conditions the disulfide bond bridging the SP and SRCR4 domains is broken, and these are visible (black arrows) below the 37 kDa and 15 kDa markers respectively. (C) Kinetics curves of NT-mini processing increasing concentrations of para-nitroaniline (pNa) conjugated peptide β (KGLVEK). Catalytic activity increases with increasing (0-10 mM) substrate concentration. Initial velocities (*V_0_*’s) were extrapolated from the 10-50 min time window. (D) Michaelis-Menten plot of *V_0_*’s versus substrate concentration. *K_m_* and *k_cat_* were estimated using non-linear regression Michaelis-Menten enzyme-kinetics models in Prism 6. Error bars represent the standard deviation of the averaged triplicate data points.

The same approach allowed for the production and purification of the inactivated Ser825Ala variant of NT-mini: NT*-mini. This mutant behaved indistinguishably form its wild-type (wt) counterpart during the production and purification processes. Predictably, no noticeable differences in yield or construct stability were observed between the two variants.

In order to determine NT-mini’s catalytic competence, its activity was investigated using synthetic chromogenic peptides and recombinantly-produced agrin constructs.

### NT-mini cleaves synthetic peptides with high specificity

Biochemical characterization of NT-mini was performed using a synthetic 6 amino acids (a.a.) peptide (β-peptide), mimicking agrin’s β-cleavage site, bearing a C-terminal chromogenic p-nitroaniline (pNa) moiety. Thus, it was possible to obtain kinetics curves by following pNa release, measured at 405 nm, over time (Fig. 1C). Initial reaction velocities (*V_0_*) extrapolated from the linear regions of those plots were used to investigate reaction parameters, compare enzymatic activities and determine kinetics. NT-mini product curves, analysed over a 1-20 mM substrate range, evidenced low affinity and slow reaction speeds. These observations were confirmed by the enzyme kinetic constants extrapolated via non-linear regression, from a *V_0_* versus substrate concentration plot (Fig. 1D). For this β-peptide, we obtained a catalytic efficiency (calculated as *k_cat_/K_m_*) of 12.9 M^−1^ s^−1^.

Contrary to these observations, previous reports indicated an inability of NT to process synthetic peptides. The absence of several NT accessory domains from NT-mini raised the question as to whether this construct might have lost the native protein’s high selectivity. Therefore, to investigate the specificity of NT-mini, we assayed its activity on a small synthetic substrate library (Suppl. Table 2), including single a.a., shorter β-peptides (5-3 mers) and also a 6 a.a. peptide mimicking the agrin α-cleavage site (α-peptide). Respective *V_0_*s, at a fixed substrate concentration (1 mM), were normalized to the reference (6-mer) β-peptide and compared (Table 1). These experiments highlighted a differential response from NT-mini dependant on the nature of the processed substrate. The α-peptide, single Lys-pNa and 3-mer β-peptide correlated with an almost complete loss of activity (−96.5, −99.5 and - 87.8%, respectively), while the 4-mer (β (4mer)) showed little variation (+7%) as compared to the 6-mer standard. Conversely, the 5-mer β-peptide substrate showed an entirely different trend, and correlated to an increased level of activity (+87%). Together, these data indicate that NT-mini is capable of catalytic activity while retaining a significant degree of substrate selectivity.

**Table 1:** Cleavage of peptide substrates by NT-mini.

| | Peptide $\alpha$ | Peptide $\beta$ | $\beta$ (5 mer) | $\beta$ (4 mer) | $\beta$ (3 mer) | Lys-pNA |
| --- | --- | --- | --- | --- | --- | --- |
| **% Activitiy** | 3.55 | 100 | 187 | 107 | 12.2 | 0.48 |
| **Difference** | -96.5 | 0 | +87 | +7 | -87.8 | -99.5 |

### NT-mini cleaves agrin-like β substrates

The initial assays carried out using synthetic peptides allowed us to investigate NT-mini’s activity, and observe catalytic competence on substrates bearing the β-cleavage site sequence. However, short peptides are a far cry from the highly glycosylated, multi-domain agrin substrate, even more so when considering its multiple splice variants. Therefore, it was mandatory to confirm that NT-mini retained the ability to cleave more native-like substrates. To this end we performed time course digestion experiments with several human agrin C-terminal constructs spanning its last three LG-EGF-LG domains (Pro1635-Pro2067) (Fig. 2A). Additionally, these recombinant substrates were designed to not only include the β-cleavage site, but also agrin’s y and z splicing insertion sites. This allowed us to address NT-mini’s β-site catalytic activity using a library of recombinant protein constructs encompassing all possible y and/or z splicing combinations (Fig. 2B). Digestion assays were performed as described in materials and methods. The reactions were analysed with SDS-PAGE, separating substrates (S) from products (P1 and P2) to generate densitometric time-resolved activity plots (Fig. 2C). Pseudo first-order rate constants for substrate consumption (*k_S_*) and LG3 (P2) product formation (*k_P2_*) were estimated via non-linear regression (Fig. 2D). *k_S_* values were then used to compare the relative activity observed for the different y and z combinations (Fig. 2E).

**Figure 2 -.**
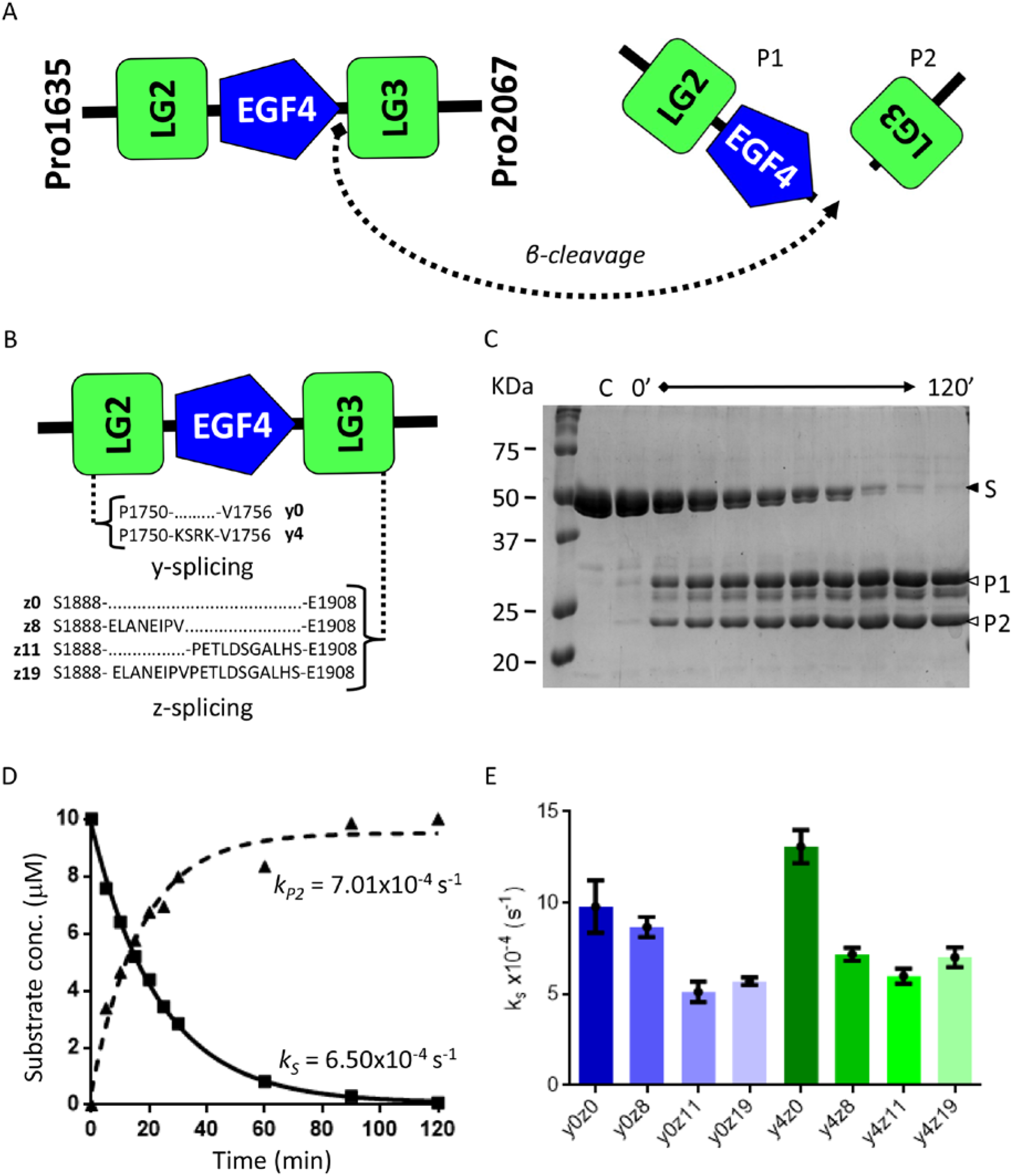
Cleavage of agrin-like substrates by NT-mini. (A) Diagram depicting cleavage of agrin-like substrates by NT-mini, resulting in the generation of two smaller product fragments corresponding to the LG2-EGF4 cluster (P1) and LG3 domain alone (P2). (B) Schematic representation of y and z splicing inserts on agrin-like substrates. (C) Representative SDS-PAGE of a time-course digestion of an agrin-like substrate with NT-mini. The substrate, S, is processed over time (120’ total) to generate two digestion products, P1 and P2. Substrate consumption and product generation are assessed by densiometric analysis. (D) Representative time resolved activity plot of product consumption and substrate generation. Curves are obtained via non-linear regression with the single phase decay (substrate) or association (product) models. These provide corresponding pseudo first order rate constants for substrate consumption (*k_S_*) and product generation (*k_P_*). (E) Comparison of *k_S_* activity rates of NT-mini on agrin-like substrates covering all splice variant combinations. Error bars represent the standard deviations.

Interestingly, NT-mini displayed catalytic competence for all assayed Agrin-like substrates with *k_S_* values ranging from 5.1×10^−4^ to 13.1×10^−4^ s^−1^ (Table 2). Surprisingly, a substrate-dependant modulation of activity was observed for the different splice variants, loosely correlating a reduction in activity to the presence of the longer z inserts (Fig. 2D). Agrin-like substrates without any z insert were among those best processed, whereas the lowest levels of activity were observed for substrates including the z11 and z19 inserts (Fig. 2D). The presence of the y4 insert further complicates the picture: the y4z0 sequence correlated to the highest levels of activity, even when compared to the splice-less y0z0 variant (Fig; 2D). However, in the presence of z inserts the effects of the y splicing site yielded cleavage rates generally comparable to their y0 counterparts. Conversely, this was not the case of the z8 variant, where the presence of the y4 insert seemed to correlate with a mild drop in activity (Fig. 2E).

**Table 2:** Cleavage rates of agrin-like substrates by NT-mini.

| Substrate | $k_S$<br>($\times 10^{-4} s^{-1}$) | SD |
| --- | --- | --- |
| *y0z0* | 9.80 | $\pm 1.43$ |
| *y0z8* | 8.68 | $\pm 0.56$ |
| *y0z11* | 5.12 | $\pm 0.56$ |
| *y0z19* | 5.70 | $\pm 0.21$ |
| *y4z0* | 13.1 | $\pm 0.91$ |
| *y4z8* | 7.19 | $\pm 0.35$ |
| *y4z11* | 5.99 | $\pm 0.42$ |
| *y4z19* | 7.02 | $\pm 0.55$ |

### Modulation of catalytic activity by metal ions

In addition to an uncanny degree of specificity, literature describes two key factors influencing NT activity, namely calcium availability^19^ and the presence (or absence) of heparin/heparan-sulphates^20^. In an attempt to identify possible domain-specific contributions to the modulation of NT activity, we investigated NT-mini’s behaviour in relation to calcium and heparin. Variations in activity were assayed using the β-peptide 6-mer synthetic substrate described previously, while alterations in protein stability were investigated with thermal denaturation experiments based on differential scanning fluorimetry (nano-DSF) techniques (Fig. 3–4).

**Figure 3 -.**
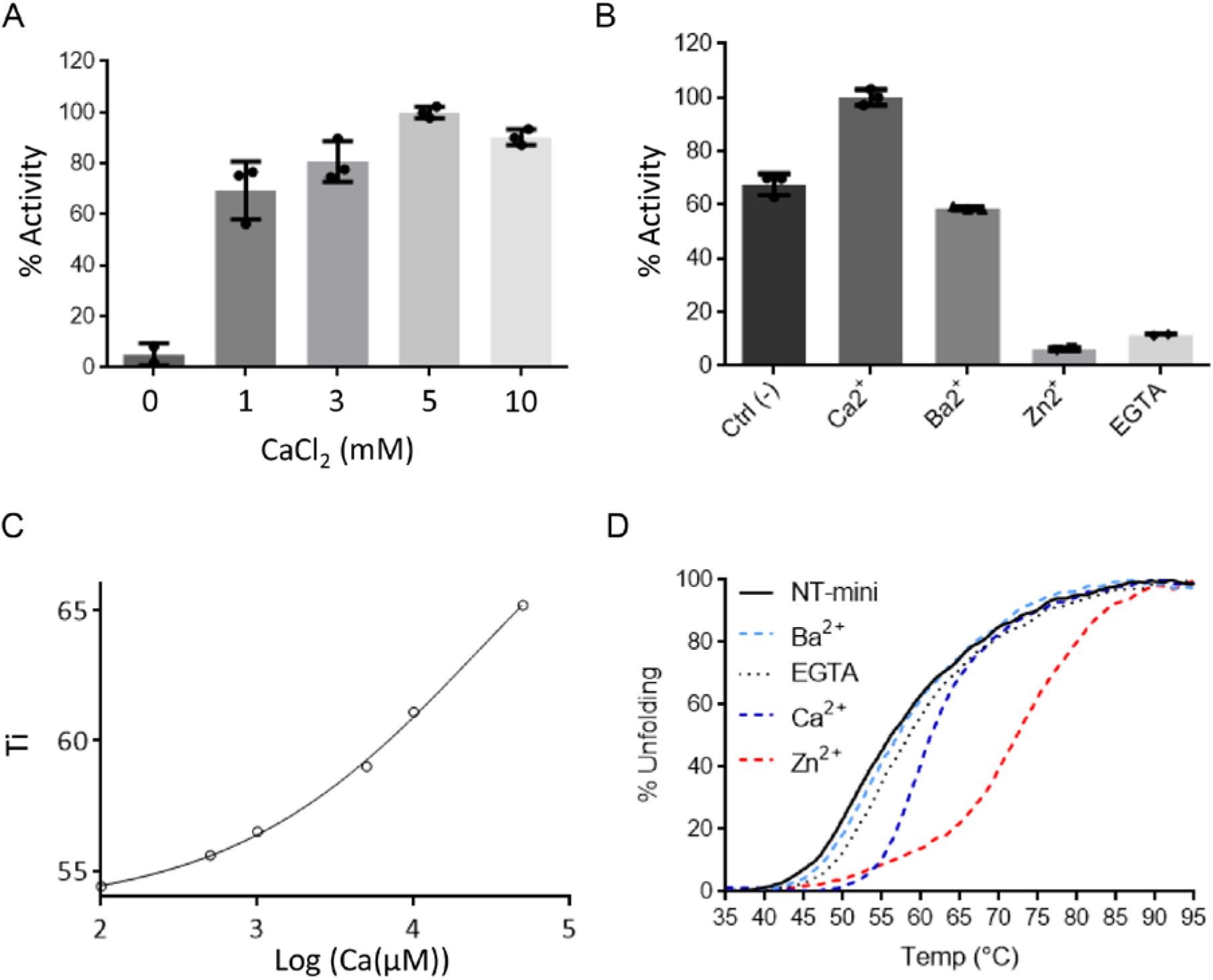
Modulation of NT-mini by bivalent metal ions. (A) Plot of NT-mini activity in presence of increasing calcium concentrations. % activity represents the NT-mini *V_0_*, extrapolated for each CaCl_2_ concentration, normalized to the highest *V_0_* value (5 mM CaCl_2_). EDTA (10 mM) was used to obtain the no-calcium reference reaction at 0 mM CaCl_2_. (B) Activity of NT-mini before the addition of metal ions, or after the addition of CaCl_2_ (5 mM), BaCl_2_ (5 mM), or ZnCl_2_ (0.2 mM). % activities represent *V_0_*’s normalized to the initial velocity of NT-mini with 5 mM CaCl_2_. (C) Thermal stabilization plot of NT-mini unfolding temperature in presence of increasing CaCl_2_ concentrations. (D) Normalized (to the highest and lowest values of each curve) plot NT-mini nano-DSF thermal denaturation curves with different bivalent metal ions. The reference curve (black line), obtained in absence of any additive, is compared to curves generated by denaturation of NT-mini in presence of 10 mM BaCl_2_ (dashed cyan line), EGTA (dotted black line), CaCl_2_ (dashed blue line) and ZnCl_2_ (dashed red line). Zn^2+^ induces the greatest degree of stabilization, Ca^2+^ has a significant but lower impact, and Ba^2+^ and EGTA have no visible effect on protein stability. Error bars on (A) and (B) indicate the standard deviation of averaged triplicate experiments.

**Figure 4 -.**
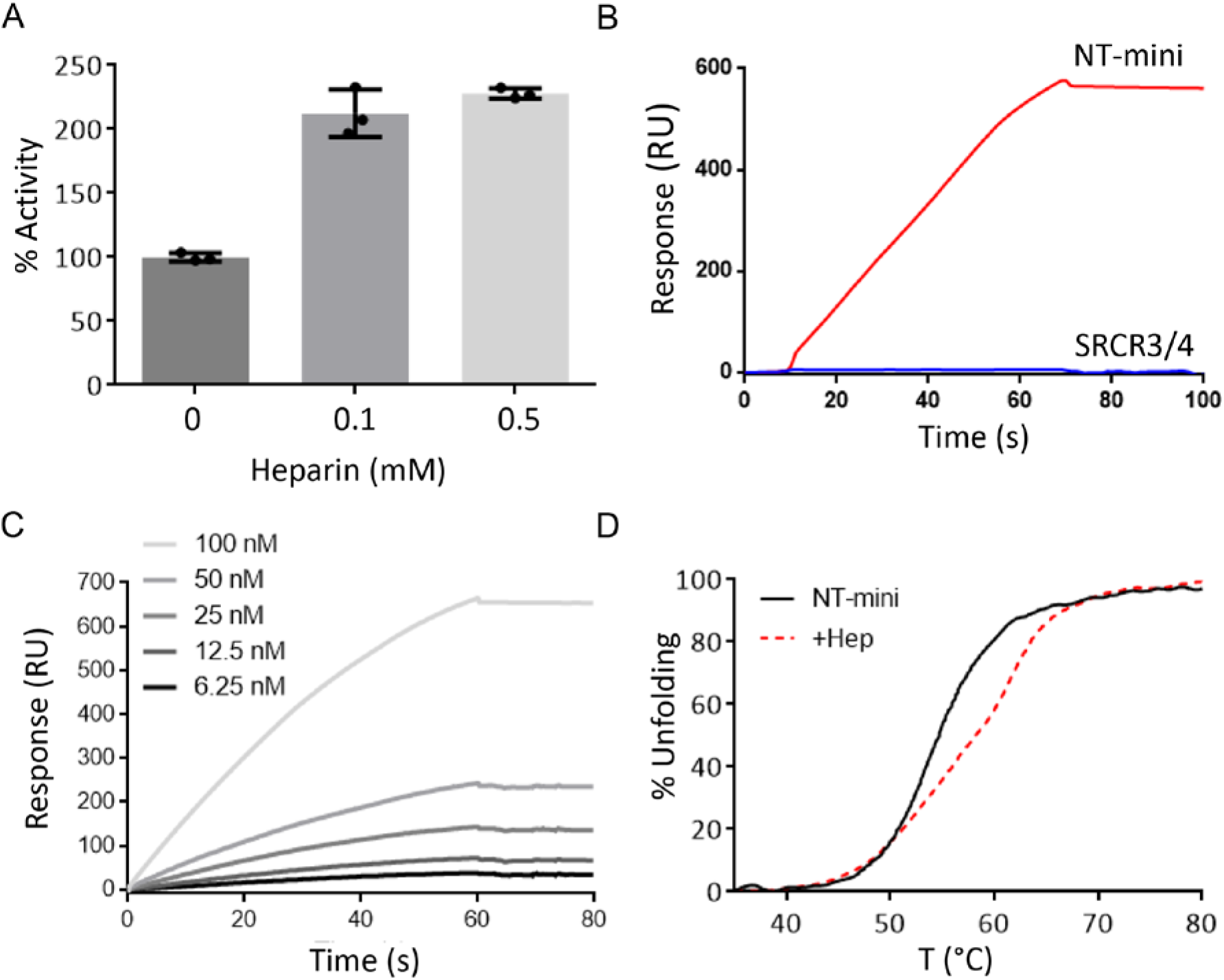
Modulation of NT-mini by heparin. (A) Activity of NT-mini in presence of heparin (0, 0.1 and 0.5 mM). % activity represents the *V_0_*’s of the reactions normalised to the reference 0 mM heparin (100%). (B) Identification of the heparin binding domain performing SPR on a heparin chip. NT-mini (red line) strongly binds the heparin chip, while the SRCR domain (blue line) preceding the SP domain does not. (C) Plot of the SPR response of increasing concentrations of NT-mini (100-6.25 nM) on a heparin chip. (D) Comparison of nano-DSF thermal unfolding of NT-mini without heparin (black line), and in presence of 1 mM heparin (dashed red line). Unfolding profiles were normalised to the highest (100%) and lowest (0%) values of each curve and used to indicate the degree of thermal denaturation (% unfolding). Error bars indicate the standard deviation of averaged triplicate experiments.

To determine if NT-mini is calcium-regulated, its catalytic activity was assessed based on the variation of β-peptide processing in relation to increasing CaCl_2_ concentrations (0-10 mM). The resulting *V_0_*’s, normalized to the highest activity, were used to compare relative reaction speeds across the assayed conditions (Fig. 3A).

NT-mini responded positively to calcium, and reaction velocities increased proportionally with the CaCl_2_ concentration. Maximal reaction velocities were reached at 5 mM, with higher concentrations yielding neither additional increases in activity nor adverse effects. This response was likely driven by high affinity interactions, as 1 mM CaCl_2_ was sufficient to elicit 70% of the observed peak activity (Fig. 3A). Additional evidence for this high specificity was found when replacing calcium with barium. Notably, at 5 mM BaCl_2_ had no significant impact on NT-mini’s activity (Fig. 3B). In light of these results, the stability of NT-mini in presence of either calcium or barium was investigated by nano-DSF. Increasing concentrations of CaCl_2_ correlated to an increasingly thermally stable protein (Fig. 3C), indicating that calcium’s role in NT activity likely extends beyond that of a “simple” co-factor. On the other hand, BaCl_2_ had no significant influence on NT-mini’s stability (Fig. 3D). Surprisingly, the effects of a different metal-ion, zinc, yielded significantly different responses. While nano-DSF experiments highlighted a beneficial impact for the protein’s stability (Fig. 3D), this was not the case for proteolytic activity. Indeed, concentrations of ZnCl_2_ as low as 0.2 mM fully inactivated NT-mini, blocking the processing of the synthetic β-peptides (Fig. 3B) and agrin-like substrates (Fig. 3B, Suppl. Fig 1). Interestingly, metal-ion chelation by EGTA resulted in comparable levels of inactivation while having no significant effect on protein stability (Fig. 3B & 3D), suggesting that zinc might have more nuanced (modulatory) implications for NT’s activity.

### Modulation of activity by heparin and identification of the responsible domain

One of the more distinctive features of NT is its reported activity modulation by heparin^20^. However, the domain(s) responsible for this property are yet to be identified. Uncertainty remains as to whether the major contributions arise from the catalytic SP domain or from the Kr-SRCR3/4 accessory domains. Therefore, our truncated constructs represented a unique opportunity to better pinpoint the predominant heparin binding domain.

NT-mini’s activity was tested for heparin modulation by carrying out in-vitro β-peptide digestion assays in presence or absence of heparin. The respective initial velocities were calculated as relative activity normalized to the reference condition without heparin (100% activity). NT-mini was seen to respond positively to heparin, and addition of 0.1 or 0.5 mM heparin induced an approximate two-fold increase in catalytic activity (Fig. 4A). As such NT’s heparin binding site was likely to be located in the two domains present in our construct. Previously,^23^ we reported the successful production and purification of the isolated SRCR domain included in NT-mini; here we used both these constructs to identify the NT domain responsible for heparin binding. Surface Plasmon Resonance (SPR) experiments using a heparin chip highlighted a clear binding response for NT-mini which was absent for the SRCR domain alone (Fig. 4B). Additionally, by assaying the SPR response to increasing NT-mini concentrations (6.25-100 nM), we could estimate the binding affinity to 19.3 pM (Fig. 5C). Interestingly, while strongly influencing activity, the high affinity interaction between heparin and NT-mini was seen to have little influence on protein stability (Fig. 4D).

**Figure 5 -.**
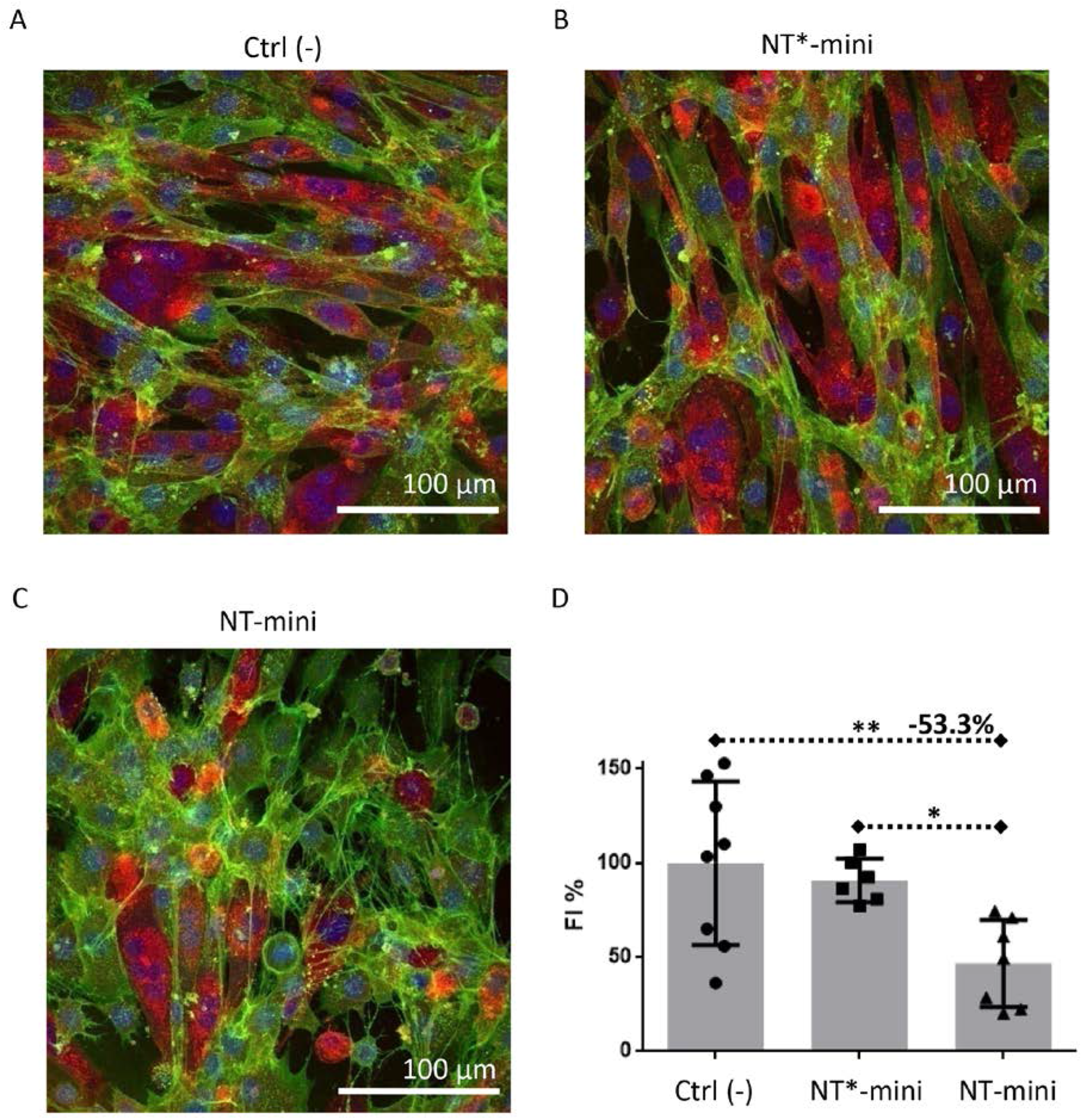
Effects of NT-mini on myotube formation. (A) Representative images of C2C12 cultures differentiated in absence of NT-mini or NT*-mini, (B) in presence of 50 ng/ml NT*-mini or (C) 50 ng/ml NT-mini. All cell images are maximum intensity Z-stack projections generated with ImageJ. (D) Comparative plot of fusion indexes normalized to non-treated controls (A, Ctrl (-)). Treatment with NT-mini hampers myotube formation causing an approximate 53% reduction in myoblast fusion, as compared to non-treated controls and NT*-mini treated cultures. Error bars indicate the standard deviation, statistical significance was determined using a one-way ANOVA coupled to a Bonferroni multiple comparison test. All analyses and plots were performed with graphpad (Prism6). *P<0.05, **P<0.01.

### Biochemical characterization of NT*-mini

Produced and purified alongside NT-mini, the inactivated Ser825Ala mutant of NT-mini (NT*-mini) was developed as a negative control for cell-based experiments. The inactivity of this construct was confirmed with the same procedures used to assay the activity of the catalytically competent NT-mini. As expected, the Ser-825-Ala mutation successfully abolished NT-mini’s proteolytic activity (Suppl. fig. 1), and NT*-mini was unable to process both the synthetic β-peptides (Suppl. Fig. 2A) and the agrin-like substrates (Suppl. Fig. 2B).

### NT-mini hampers myotube formation in-vitro

Based on our *in-vitro* biochemical characterization, NT-mini seemed to recapitulate the properties of the full-length protease. Coupled with the inactivated NT*-mini, they represented a unique set of tools to try and unravel the biological function of NT. Thus, we wanted to establish whether NT-mini could mimic its parent protein in a cellular environment. To this end, we evaluated the impact of NT-mini on myotube formation^11^ using C2C12 mouse myoblasts, a subclone of a well-documented cell line (C2) that is often used in studies addressing muscle cell development and NMJ maturation^26–32^. Myoblast differentiation was induced by serum starvation protracted for a total of 7 days. Throughout this time-period the differentiating cells were left untreated (controls), or treated with either NT-mini or NT*mini. Fully “mature” cultures were stained to detect cell nuclei (blue), acetylcholine receptors (AChRs, red), and cell-surface glycans (green) (Suppl. Fig. 3A). Cell boundaries were determined based on the glycan and AChR stains, and differentiated myotubes identified as cells containing 3 or more nuclei (Suppl. Fig. 3B-C). The effect of NT-mini’s activity on myotube formation was then quantified and cross-compared to NT*-mini and the control based on the relative fusion indexes (FI), corresponding to the respective ratios of nuclei incorporated in myotubes to the total number of nuclei.

Treatments with NT-mini had no visible impact on AChR distribution, which appeared generally diffuse throughout myotube membranes regardless of treatments. Rare, sporadic, small clusters were occasionally observed, but seemed to bear no relation to the addition of NT-mini, and were also found in presence of NT*-mini as well as non-treated controls. However, when myotube formation was assessed based on the relative FIs, NT-mini was seen to have a negative impact on the process. While the inactive NT*-mini showed no significant degree of variation compared to non-treated controls, NT-mini’s activity was seen to correlate to a significant (≈53.3 %) drop in FI (Fig. 5), indicating a reduced degree of myotube formation. This seemed to reflect the reported muscle fibre loss in NT overexpressing mice^11^, and hints at a more direct involvement of NT in the process than might have been previously supposed.

## Discussion

Proteases are known to play essential roles in processes of synapse formation and rearrangement, both in the CNS and at peripheral synapses like NMJs^5^. In this regard, the NT-agrin enzyme-substrate pair represent a signalling pathway, associated to synaptic rearrangement, which is shared across central and peripheral synapses. Agrin has long been known to play a direct role in synaptic organisation^10,33–39^ however the contributions of NT to the same processes (agrin-dependant or otherwise) are generally less understood. While a number of observations have tentatively associated NT activity to synaptic plasticity^10,11,15,40,41^, a more detailed molecular understanding of NT functionality and its interplay with agrin is lacking. The difficult recombinant production of full-length NT has been at the core of this limitation^19^. In this context, our shorter construct (NT-mini) including its SP and most proximal SRCR domain and its inactive variant NT*-mini are more amenable to recombinant production using suspension-cultured HEK293 cells with reasonably good protein yields (≈ 1 mg/L of culture), and represent novel tools to better understand the molecular implications of NT activity and its regulatory mechanisms.

Despite its truncated nature, NT-mini conserved catalytic activity, and was capable of processing the β cleavage site when presented on both agrin-like substrates and, with much lower efficiency (the affinity for β-like peptide substrates falls in the mM range), synthetic peptides of different lengths. Out of several peptide-substrates, the single lysine amino acid and short 3-mer peptide substrates were the only ones to be virtually unprocessed. This is indicative of the conservation of NT’s high specificity in NT-mini, especially in light of the successful processing of the longer β-like peptides. In this context the 5-mer peptide substrate (correlated to the highest cleavage rates) is particularly interesting, as it seems to highlight a sort of catalytic “sweet-spot”. This indicates that the corresponding amino acids (-GLVEK-) might represent a preferred substrate recognition sequence, or that longer polypeptides may require additional long-range binding sites for optimal processing by the enzyme.

Surprisingly, NT-mini’s response to the α peptide was significantly different, as we were unable to observe any significant processing. It is possible that the lack of activity on that substrate is representative both of high specificity, and of non-permissive conditions for α-site cleavage. This might suggest that NT-mediated cleavage of agrin at that site could be subordinate to exosite interactions extending beyond NT’s SP domain and the recognised α-site sequence. These are likely to include the accessory domains absent from NT-mini, as well as ulterior agrin domains and the post-translational modifications (such as heparan sulphate side-chains) proximal to the α cleavage site.

Collectively, this seems to indicate that the missing NT accessory domains are not strictly necessary for catalytic competence, at least insofar as β-site cleavage is concerned. Interestingly, these observations not only highlight the conservation of NT’s high specificity in the much shorter NT-mini, but also provide a possible insight into the function of NT’s accessory domains as it has long been speculated that these might find a role in mediating protein-protein interactions such as substrate recognition. Accordingly, the absence of several of these domains from NT-mini seems to have influenced its specificity, allowing it to process short peptide fragments. As such, it seems plausible that NT’s specificity might be significantly influenced by its accessory domains, and that its full activity on agrin is dependent as much on cleavage site recognition as it is on NT-agrin exosite interactions extending beyond the SP domain. This could offer a possible explanation as to why NT-mini fails to recognise and cleave α-site mimicking peptides. Indeed, in such a setup all interactions beyond those involving the active site and the cleaved sequence are likely absent, something which would negatively impact any catalytic activity reliant upon broader interactions stretching across other domains.

Regardless of the low affinities involved, NT-mini’s processing of β-like peptides allowed us to investigate its activity in a range of conditions, and compare its behaviour to that of wild-type NT^19,20^. This highlighted that NT-mini’s activity is both calcium dependent and heparin modulated, behaviours which reflect what has been previously reported regarding murine NT. Furthermore, given the truncated nature of NT-mini, these observations indicate the presence of calcium and heparin binding sites within the final two domains of this protease. This is especially true regarding heparin, as it was possible to determine that binding with picomolar affinity occurs only in presence of the SP domain.

Alongside these considerations, the previously reported calcium binding properties of the murine NT SRCR3 domain^23^ seem to indicate that calcium binding by NT is likely to occur at multiple sites throughout the protein. This seems even more plausible when taking into account the high degree of structural conservation found across the SRCR domain class^23,42–44^. Interestingly, NT-mini itself is likely to contain multiple binding sites, as its interaction with calcium can be associated to two effects observable across distinct concentration ranges. Namely, while NT-mini reaches peak catalytic activity in presence of 5 mM CaCl_2_, this is not the case for its calcium-induced thermal stabilisation which continues to improve with increasingly higher CaCl_2_ concentrations. Therefore, it would be plausible to associate the calcium-dependent activity to a binding site on the SP domain, while the observed calcium-induced stabilization likely stems from binding at both the SP and SRCR domains.

Investigating the specificity of the interaction yielded novel insights into the response of NT to another bivalent metal ion essential to the CNS. Initial experiments with barium seemed to indicate that NT-mini’s interactions with metal ions was strongly calcium specific. However, when assaying its response to zinc NT-mini showed an unexpected behaviour. On the one hand Zn^2+^ ions were seen to be strongly beneficial for the protein’s stability (more so than calcium), while on the other their presence was highly detrimental for its catalytic activity. This would seem to hint at a dual regulation of NT-mini (and therefore NT) by different bivalent ions. Calcium would serve to stimulate catalytic activity, while zinc might be responsible for its down-regulation. Given the significant importance of Ca^2+^ and Zn^2+^ currents in synaptic signalling within the CNS^45–47^, it would be interesting to speculate that NT might be capable of responding *in-vivo* to the different stimuli. This would, in turn, allow spatio-temporal fine-tuning of NT’s activity not only on the basis of neuronal activity, but also in relation to the fluctuations in zinc/calcium currents. Something which would be very much in agreement with the documented importance of NT activity for CNS/PNS synaptic plasticity.

Zinc, calcium and heparin are not, however, the only factors that seem to modulate NT-mini’s activity. Indeed, when testing its catalytic competence on short C-terminal recombinant fragments of agrin, NT-mini was seen to have a substrate modulated activity dependant on the agrin splicing variations present on the substrate. This might represent another modulatory node possibly partially independent from NT itself but instead strongly tied to the relative abundance of agrin splice variants. Their different cleavage rates might result in the accumulation of different agrin fragments, with potentially different biological roles, in a manner directly dependant on the production, distribution, and presence of substrate variants. In turn, that would lead to interesting modulatory possibilities that might account for Agrin’s multiple reported biological roles and represent a means of spatio-temporally regulating agrin-mediated signalling.

Given the modulable nature of NT enzymatic activity, its effect on Agrin signalling, and its involvement in processes of synaptic reorganisation, it is surprising that deregulation of the NT-agrin axis has been linked only to a small number of pathological states^11,40,41,48^. One of the few known pathogenic implications of NT deregulation is centred on the formation and maintenance of NMJ’s. Overexpression, and thus hyperactivity, of NT in mice has been documented to cause an acceleration in the NMJ “life-cycle” leading to a premature degradation correlated with generalized muscular weakness and wasting^11^. However, the highly interconnected nature of these synapses, makes it difficult to say whether this phenotype is directly caused by excessive NT activity. It might be subordinated to a reduction in muscle contraction caused by impaired NMJ signalling. Our experiments performed with NT-mini on C2C12 myoblast differentiation offer an insight in this process. The activity of NT-mini was directly responsible for a reduction in myotube formation, something which reflects the muscle wasting observed in NT overexpressing mice. Therefore, it is possible that the muscle weakness/wasting observed in the mouse model could be more strongly dependant on NT’s activity than might have originally been thought.

In conclusion, it would seem that NT-mini may offer an effective means of expanding the molecular understanding of NT. This is especially true as NT-mini’s activity seems to respond to the same modulators affecting full-length NT. Furthermore, the characterisation of this truncated NT variant has allowed not only to identify the NT heparin binding domain, but has also brought light on previously undescribed features of NT. Namely, that its activity seems to be subject to additional substrate-dependant and zinc-dependant regulation. Together, this data highlights how the NT-agrin signalling axis might be fine-tuned by both the alternative splicing of the substrate and the Ca^2+^/Zn^2+^ ion currents present at synaptic interfaces. Something which appears consistent with the documented contributions of both NT and agrin to CNS and PNS synaptic plasticity. Finally, given the similarity of its activity to NT, NT-mini represents a unique tool to investigate NT activity in “controlled” cell-based experiments. If tested together with its inactivated mutated variant NT*-mini, it might allow to unravel the possible cellular implications of NT’s activity. Something which would be the first critical step towards a better understanding of its role in a broader and complex biological context.

## Materials and methods

### Molecular cloning

Recombinant constructs for NT-mini (Gly497-Leu875) and C-terminal Agrin substrates (Pro1635-Pro2067) were amplified using a Phusion DNA polymerase (Thermo Fisher Scientific) starting from a source synthetic gene (GeneWiz) corresponding the full-length human NT cDNA (UniProt P56730) and from a human Agrin y0z0 cDNA (UniProt O00468) obtained from Source Biosciences, respectively, using the oligonucleotides listed in Suppl. Table 1.

The resulting PCR products were subcloned into pCR4-TOPO vectors (Thermo Fisher Scientific) using BamHI and NotI restriction sites. These plasmids served as the basis for the generation of the inactivated Ser-825-Ala NT*-mini and of the various Agrin splice variants via site directed mutagenesis. The corresponding PCR reactions were performed with Phusion DNA polymerase (Thermo Fisher Scientific) using the oligonucleotides listed in Suppl. Table 1.

All constructs were verified using Sanger sequencing (Microsynth) and then were sub-cloned into modified pUPE.106.08 plasmids (U-Protein Express BV) for secreted protein expression in mammalian cell culture systems. For NT-mini, the final expression constructs contained an N-terminal 8xHis-SUMO tag used to stimulate expression and facilitate purification. For Agrin constructs, the final expression construct contained an N-terminal 6xHis-tag followed by a Tobacco Etch Virus Protease (TEV) cleavage site.

### Recombinant protein expression

All NT and Agrin constructs were produced via transient transfection of SFM-HEK293 (Expression Systems) cell suspension cultures adapted to grow in a serum-free medium (ESF, Expression Systems). Transfections were performed using linear polyethylenimine (PEI MAX, Polysciences) in a 5:1 (w:w) PEI:DNA ratio. Zymogen-to-enzyme activation of NT-mini constructs was performed during protein production by co-transfecting the NT-mini plasmid in a 1:5 ratio with a pUPE expression plasmid for mammalian expression carrying the human form of the pro-protein convertase Furin cDNA (U-Protein Express BV). Four hours post-transfection, the culture was supplemented with peptone supplement (Primatone RL, Sigma-Aldrich) at a final concentration of 0.6% (w/v). Transfected cultures were maintained for 6 days to allow recombinant protein expression and accumulation. On the 7^th^ day the growth media was harvested by centrifugation. Pellets were discarded and the recombinant proteins were purified from the supernatant using chromatographic approaches.

### Recombinant protein purification

Cell media were adjusted with concentrated buffer to a final 200 mM NaCl, 25 mM HEPES/NaOH, pH 8 and filtered through a 0.45 μm syringe filter (Sarstedt) using a peristaltic pump (Thermo Fisher Scientific). All chromatographic steps were carried out using an NGC chromatography system (Biorad), by monitoring UV absorbance at 280 nm. Samples from each step were further analysed with reducing and non-reducing SDS-PAGE. Purification of NT-mini or NT*-mini was obtained with a three-step Heparin/Ni-IMAC “mixed-mode” approach, and completed by size exclusion chromatography (SEC). The filtered medium was supplemented 5 mM CaCl_2_ and loaded, at room temp, on a 20 mL Heparin column (GE Healthcare) pre-conditioned with buffer NT.A (200mM NaCl, 5 mM CaCl_2_, 25 mM HEPES/NaOH, pH 8). Unbound material was washed off the column with buffer NT.A, and weakly bound contaminants were removed by increasing the NaCl concentration to 250 mM. The protein of interest (NT-mini or NT*-mini) was then eluted into a 5 ml Ni-IMAC HisTrap Excel column (GE Healthcare) by further increasing the NaCl concentration to 600 mM (buffer NT.B). Unbound material was washed off the Ni-IMAC column with buffer NT.A, after which the recombinant protein was eluted using buffer NT.C (200mM NaCl, 5 mM CaCl_2_, 250 mM imidazole, 25 mM HEPES/NaOH pH 8) into a 1 mL HiTrap Heparin HP column (GE Healthcare). This column was further washed with buffer NT.A, and the bound NT construct was de-tagged by loading the column with 10 mL of a 4 μg/mL SUMO protease solution. Tag cleavage was allowed to proceed in column overnight at 4 °C, then the cleaved tag was washed off the column with buffer NT.A. At this point NT-mini/NT*-mini was eluted from the heparin column using buffer NT.B. The obtained material was concentrated to reach a final volume ≤0.5 mL and further purified by SEC on a Superdex 200 10/300 Increase column (GE Healthcare), equilibrated with buffer GF (200 mM NaCl, 25 mM HEPES/NaOH, pH 8). Finally, the purified NT-mini/NT*-mini was concentrated to 1 mg/mL with a Vivaspin Turbo10 kDa MWCO concentrator (Sartorius), flash frozen in liquid nitrogen and stored at −80 °C. This material served as the stock solution for all subsequent *in-vitro* assays and cell culture experiments.

For Agrin constructs, the filtered medium was loaded on a 5 mL HisTrap Excel column (GE Healthcare) equilibrated with buffer AG.A (500mM NaCl, 25 mM HEPES/NaOH, pH 8). Non-specific contaminants were removed by washing the column with buffer AG.A supplemented with 25 mM imidazole, then the elution of recombinant agrin constructs was obtained using buffer AG.B (500mM NaCl, 250 mM imidazole, 25 mM HEPES/NaOH, pH 8). Fractions were pooled and dialyzed overnight against buffer AG.A in the presence of 25 μg/mL His-tagged TEV protease. Tag and TEV protease removal was achieved by re-loading the dialyzed sample onto the 5 mL HisTrap Excel column (GE Healthcare), and collecting the unbound tagless sample. Fractions containing pure agrin constructs as assessed by SDS-PAGE analysis underwent concentration through a Vivaspin Turbo 30 kDa MWCO concentrator (Sartorius) to reach a final volume below 0.5 mL, and further purified by SEC on a Superdex 200 10/300 Increase column (GE Healthcare) equilibrated with buffer GF. All samples eluted as single peaks distant from the column void volume. Pooled fractions were concentrated to 3-10 mg/mL, flash frozen in liquid nitrogen and stored at −80 °C until usage.

### Activity assays with peptide substrates

NT-mini biochemical characterization was performed colorimetrically, at a fixed enzyme concentration of 1 μM, with p-nitroaniline (pNa)-bearing peptide substrates (Suppl. Table 2) allowing to follow product generation as an increase in absorbance at 405 nm.

Time-course measurements were performed at 37 °C in a clear-bottom 386 well plate (Greiner) in GF buffer. For each measurement, the final reaction volume was 20 μL. Substrate and CaCl_2_ stock solutions were prepared, respectively, as 100 mM and 1 M stocks in 25 mM HEPES/NaOH pH 8 and diluted as necessary. Stocks of BaCl_2_ and ZnCl_2_ for the corresponding experiments were prepared in identical fashion as those of CaCl_2_. Heparin stocks were prepared as a 1 mM solution by directly dissolving heparin sodium salt (Sigma #H3393) in reaction buffer. For experiments assessing activity in relation to ionic strength, the NaCl concentration of the reaction buffer was altered to cover a 100-500 mM range, reaction parameters were otherwise unchanged from reference conditions. Reference reactions were performed using 1 mM CaCl_2_ and 5 mM peptide substrate, respectively. Each reaction was prepared on ice, with 1 mg/mL NT-mini as the last component, and briefly mixed by pipetting before measurement. To prevent evaporation each sample was then overlayed with a small drop (~7 μL) of paraffin oil. Triplicate kinetics measurements were performed using a CLARIOstar plate reader (BMG LABTECH) over the span of 230 min following absorbance at wavelengths of 405 and 600 nm. Measurements at 600 nm were used to monitor and correct for the appearance of artefacts. Absorbance values were converted to concentration curves by accounting for path length and using the pNa extinction coefficient at 405 nm (9500 M^−1^ cm^−1^). The linear region of these plots (50 points in the 10-50 min measurement interval) was used to extrapolate initial reaction velocities (*V_0_*) via linear regression. The *V_0_* values were directly fitted to the Michaelis-Menten equation using Prism 6 (GraphPad), which provided the values of apparent *k_cat_* and *K_m_* along with their associated errors. Propagation of statistical error value during the calculation of *k_cat_*/*K_m_* values was carried out as described^49^. Absence of proteolytic activity in NT*-mini was assessed in the same manner.

### Time course digestions of Agrin-like substrates

Catalytic competence on native agrin-like substrates was assessed via time course assays using purified recombinant human agrin LG2-LG3 constructs. Digestions with NT-mini at 5 μg/mL were performed at a fixed starting substrate concentration of 0.5 mg/mL in GF buffer supplemented with 5 mM CaCl_2_. All necessary dilutions were performed with the reference reaction buffer.

Typical reaction mixes were prepared on ice in a PCR tube (total reaction volume 110 μL) and were composed of 500 μg/mL agrin construct, 5 mM CaCl_2_, 5 μg/mL NT-mini. NT-mini was added last and then the solution was mixed by gentle pipetting. Reaction mixes were maintained at 37 °C in a PCR thermocycler (Eppendorf) and 10 μL samples were collected after 0, 5, 10, 15, 20, 25, 30, 60, 90 and 120 minutes. Each sample was immediately supplemented with 5 μL of SDS-PAGE reducing sample buffer, boiled for 10 min at 95 °C to block the reaction and loaded on a 16% polyacrylamide gel. Protein bands were stained with colloidal Coomassie blue solution (0.02% Brilliant Blue G250 *w/v*, 20% ethanol *v/v*, 5% Al2(SO4)3 *w/v*, 1.05 M phosphoric acid) overnight and carefully destained with destaining solution (10% acetic acid *v/v*, 20% ethanol *v/v*). Images for analysis were taken with a ChemiDoc imager (BioRad) and substrate/product band densities were measured with imageJ^50^. The intensity values for the substrate bands obtained after at 0 minutes of digestion were set to represent starting substrate concentration (0.5 mg/mL or 10 μM), and the normalized intensities associated to subsequent time point bands were then plotted as a function of time. Prism6 (Graphpad) was used to plot a non-linear regression curve from which pseudo first order rate constants for substrate consumption (*k_s_*) and product generation (*k_p_*) were derived, along with their associated errors. The same procedure was followed for experiments with NT*-mini.

### Thermal denaturation assays

The effects of calcium, zinc, barium, heparin, EGTA or EDTA on NT-mini’s stability were assayed directly using thermal denaturation coupled to nano-differential scanning fluorimetry (nano-DSF). For these experiments the concentration of NT-mini was fixed at 2 μM (0.1 mg/mL), while bivalent metal ions (Ca^2+^, Zn^2+^, Ba^2+^) were assayed at 5 mM, and chelators (EGTA, EDTA) were tested at 10 mM. All chemicals (Sigma-Aldrich) were dissolved directly in GF buffer, to prepare stock solutions at high concentrations: CaCl_2_ 1 M, BaCl_2_ 1 M, ZnCl_2_ 1 M, NT-mini 20 μM (≈ 1 mg/ml), EGTA 1 M and EDTA 1 M. The same buffer was also used for all necessary dilutions to reach the assay concentrations from those stocks. Thermally-driven protein unfolding was monitored in Tycho quartz capillaries (Nanotemper) using a Tycho NT.6 instrument (Nanotemper), which allowed for determination of NT-mini’s unfolding temperatures in the different assayed conditions. The smoothed data was re-plotted using Prism 6 (Graphpad) for comparative purposes.

### SPR determination of heparin binding and affinity

Measurements of heparin binding were performed in a Biacore T200 instrument (GE Healthcare) using a heparin-coated chip (Xantec). All analyses were performed in tris-buffer saline (TBS, 150 mM NaCl, 50 mM Tris-HCl pH 7.5) unless otherwise noted. After each sample injection, the chip was regenerated using a TBS solution supplemented with 1 M NaCl. For the identification of the heparin binding domain, the murine SRCR3 (human SRCR4) domain and NT-mini were run individually at a concentration of 50 nM with a contact time of 60 s at a flow of 50 μL/min. For the estimation of the NT-mini’s affinity for heparin, a decreasing series of NT-mini concentrations (25, 5, 1, 0.2 and 0.04 nM) were obtained by serial dilution in TBS, starting from a protein stock of ≈22 μM. Steady-state binding was assayed with triplicate experiments performed using the single cycle kinetics mode with a contact time of 60 s and a flow rate of 30 μL/min. Curves were analysed using the kinetics fit of the Biacore T200 evaluation software (GE Healthcare). For visualization purposes, the SPR traces were exported and re-plotted using prism 6 (Graphpad).

### C2C12 cultures

For maintenance C2C12 myoblasts (kindly provided by DM Rossi, ICS Maugeri, Pavia) were grown in 75 cm^2^ T-flasks (VWR) at 37 °C in a humidified 5% CO_2_ atmosphere. Cells were maintained in a high-glucose DMEM (Thermo Fisher Scientific) medium supplemented with 10% (*v/v*) foetal bovine serum (FBS, Thermo Fischer Scientific) and 1% (*v/v*) non-essential amino acids (NEAA, Thermo Fisher Scientific). Growth medium was refreshed on average every 48 h and cells were generally split 1/10 when at 60-70% confluence.

To perform differentiation experiments, cells were transferred to 6-well plates containing sterilized glass cover slips and seeded at ~50% confluence. Cultures were allowed to stabilize for 24 h before inducing differentiation. This was obtained using high glucose DMEM medium with a reduced FBS content (1%) but still supplemented with 2% NEAA. Differentiation was generally protracted for 7 days before proceeding with immuno-staining. Growth media was refreshed every 48 h.

Treatments with NT-mini/NT*-mini (50 ng/mL) were performed throughout the 7-day differentiation period by supplementing the differentiation medium with the purified recombinant protein. All necessary stock dilutions were performed directly in differentiation medium.

### C2C12 staining and analysis

Evaluation of myoblast-to-myotube differentiation was carried out using a live stain protocol adapted from the procedures described in Stanga et al. 2016^32^, McMorran et al. 2017^51^, and Harrison at al. 1992^52^. Briefly, cellular boundaries were identified by staining cell-surface glycans with an Alexa645-wheat germ agglutinin (WGA) conjugate, while nuclei were stained using Hoechst dye. Acetylcholine receptor (AChR) clustering was evaluated using an Alexa594-bungarotoxin (Btx) conjugate. Hoechst and Btx stains were added directly to the media in the 6 well plate, in a 1/2000 (*v/v*) and 1/1000 (v/v) ratio respectively, and incubated at 37 °C for 45’. Staining with WGA was also performed directly in the culture media, but in a 1/250 (v/v) ratio and with a 15’ incubation time. After staining, the cells were washed once with PBS, fixed with a 4% (*w/v*) paraformaldehyde (PFA) in PBS solution at room temperature for 15’, and washed again with PBS. Finally, the cover slips were mounted on microscopy slides with a drop of mounting media (Sigma-Aldrich). These mounts were wrapped in tin foil, and allowed to set O/N at 4 °C before being imaged using a SP8 white-laser confocal microscope (Leica Microsystems). Filters were adjusted specifically for each stain to visualize nuclei (Hoechst, Ex 405 nm, Em 429 nm), AChRs (Btx, Ex 598 nm, Em 634 nm) and cell-surface glycans (WGA, Ex 653 nm, Em 692 nm). Z-stacks of 0.5 μm thickness were collected at 40x magnification, and images were analysed with imageJ^50^. Myotube identification and assignment of nuclei were performed by manual slice-by-slice assessment of the Z-stacks. Fusion indexes were calculated as the fraction of total nuclei incorporated into myotubes. These data were normalised to the non-treated control conditions, plotted on Prism 6 (Graphpad), and the statistical significance of the observed differences assessed performing a one-way ANOVA with a multiple comparison (Bonferroni) test.

## Acknowledgements

We would like to thank Dr. M. Palamini and Dr. M. Campioni for support in cell culture setup, and Dr. D.M. Rossi for provision of the C2C12 cells used in this study.

## Author Contributions

AC and FF designed research. FF, PKC, and SS supervised research. AC generated and purified all recombinant proteins and carried out biochemical characterizations, with support from SF with cell cultures. CC purified NT*-mini. AC, SF and SS carried out cell-based experiments. AC and FF analysed the data and wrote the manuscript, with contributions from all authors.

