## Supplementary material for "NT-mini, a recombinant tool for the study of Neurotrypsin functionality": Suppl.

Anselmo Canciani *et al.*

**Supplementary Table 1: List of primers used for construct amplification.** Restrictions sites are underlined. Modified codon for Ser-825-Ala mutation indicated in italics.

| **Construct** | | **Primer sequence (5'-3')** |
|---|---|---|
| **NT-mini** | **Fwd** | AAAGGATCCGGTTTTCCTGTCAGACTGATGGATGG |
| | **Rev** | TAAGCGGCCGCCAGTTTGGTGACACTTTTTATCCAAGGTACAAAGG |
| **NT*-mini** | **Fwd** | *GCT*GGAGGACCACTCATGTGTGA |
| | **Rev** | GTCTCCCTGGCAGCTGTCCA |
| **Agrin LG2-LG3** | **Fwd** | AAAAGGATCCCCCTTCCTGGCTGACTTCAAC |
| | **Rev** | AAAAGCGGCCGCTGGGGTGGGGCAGGGCCG |
| **Agrin y4** | **Fwd** | CGTGTGTTGGGGGAGTCCCCGAAAAGCCGCAAAGTTCCGCACACCGTCCTCAACCTG |
| | **Rev** | CAGGTTGAGGACGGTGTGCGGAACTTTGCGGCTTTTCGGGGACTCCCCCAACACACG |
| **Agrin z8** | **Fwd** | GCCAACGAAATTCCGGTGGAGAAGGCACTGCAGAGCAA |
| | **Rev** | CCGGAATTTCGTTGGCCAGTTCGCTCTCGGTCACAGCGTTG |
| **Agrin z11** | **Fwd** | CTGGATAGCGGCGCGCTGCATAGCGAGAAGGCACTGCAGAGCAA |
| | **Rev** | CGCGCCGCTATCCAGGGTTTCCGGGCTCTCGGTCACAGCGTTG |
| **Agrin z19** | **Fwd** | TGGCCAACGAAATTCCGGTGCCGGAAACCCTGGATAGC |
| | **Rev** | GCTATCCAGGGTTTCCGGCACCGGAATTTCGTTGGCCA |

**Supplementary Table 2: synthetic substrates used for NT-mini characterization.**

| | Peptide α | Peptide β | β (5 mer) | β (4 mer) | β (3 mer) | Lys-pNa |
|---|---|---|---|---|---|---|
| **Sequence** | GPPVER-pNa | KGLVEK-pNa | GLVEK-pNa | LVEK-pNa | VEK-pNa | K-pNa |
| **Source** | *China Peptides* | | | | | *BACHEM* |

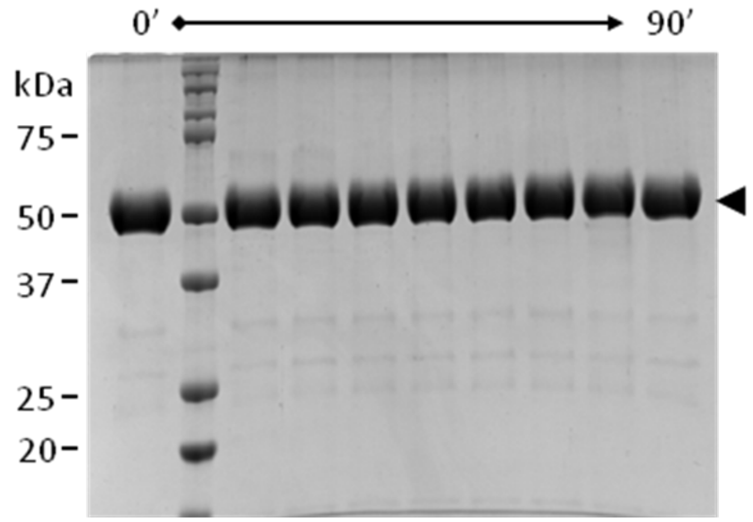


**Supplementary figure 1 - Zinc blocks digestion of agrin-like substrates by NT-mini.** SDS-PAGE of a time a resolved digestion assay of an agrin-like substrate (black arrow) by NT-mini in presence of 0.2 mM ZnCl2. The presence of zinc strongly inhibits NT-mini leaving the substrate unprocessed.

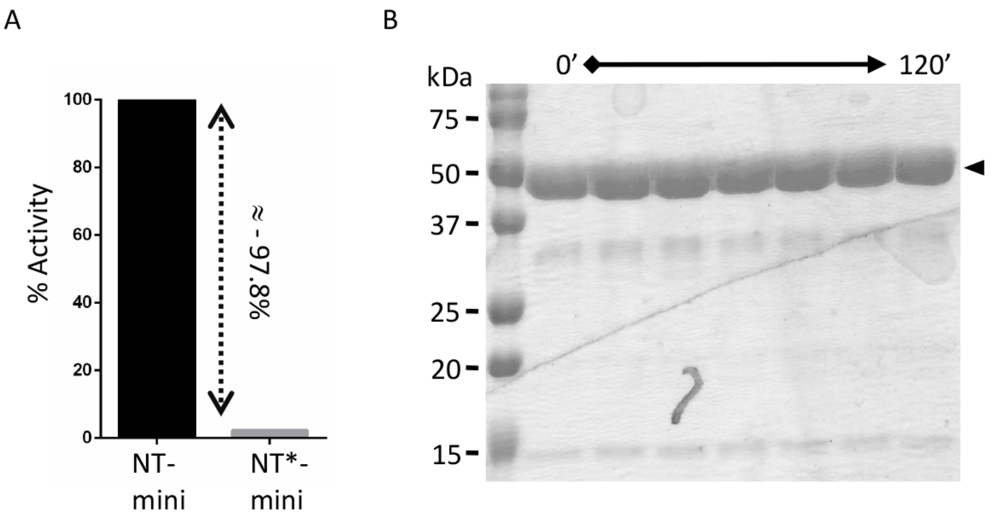


**Supplementary figure 2 - Assessment of inactivity for NT*-mini.** (A) Activity of NT*-mini on the reference synthetic peptide β as compared to NT-mini. % activity is normalized to the $V_0$ of NT-mini. (B) SDS-PAGE showing a time-course of non-digestion of an agrin-like substrate (black arrow) by NT*-mini.

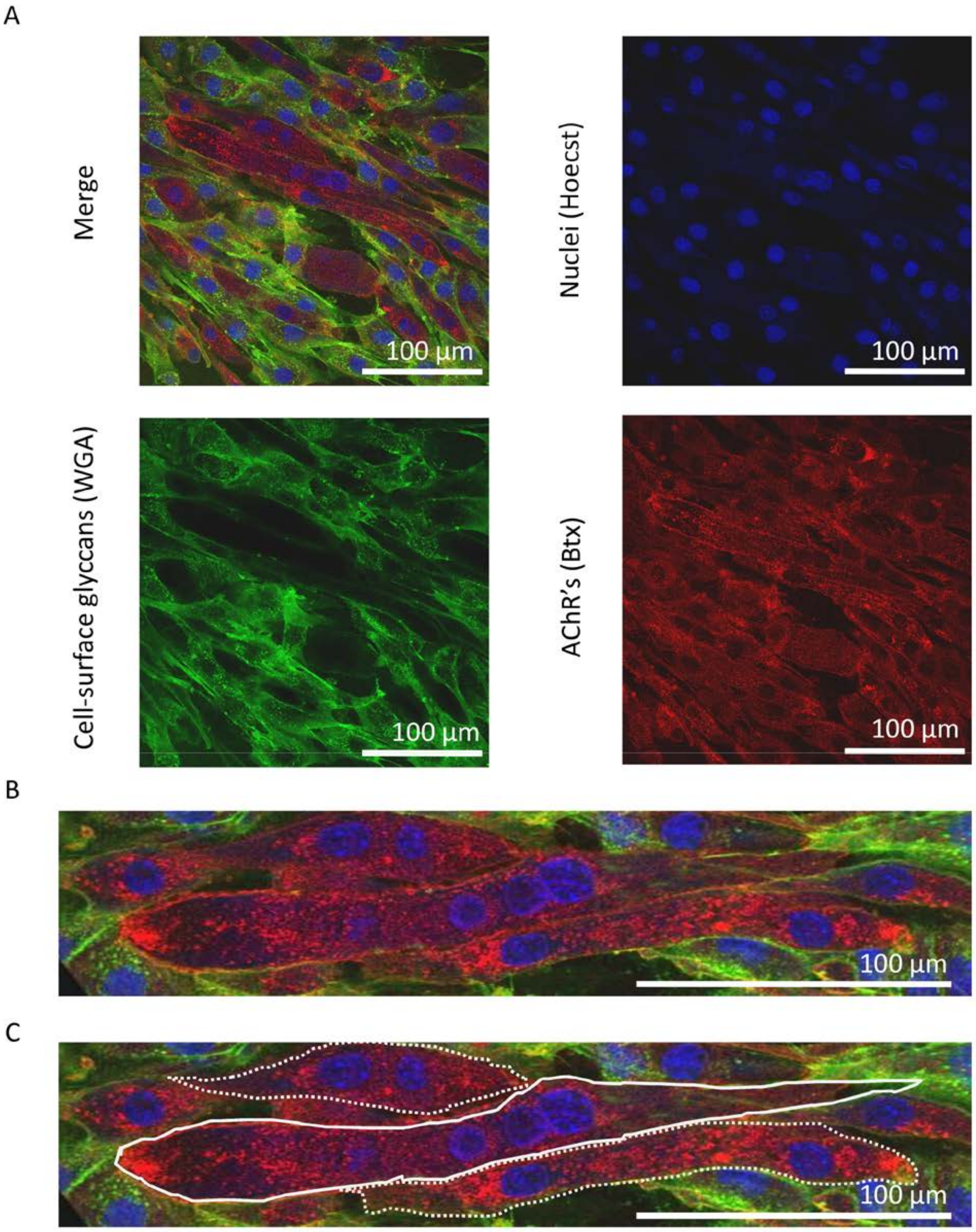


**Supplementary figure 3 - C2C12 staining and myotube identification.** (A) Single confocal Z-stack slice images, collected at 40x magnifcation, representative of myotube staining. Nuclei are stained with Hoechst (blue), cell-surface glycans are stained with a WGA-Alexa647 (wheat germ agglutinin-Alexa647) conjugate (green), and acetylcholine receptors (AChRs) are stained with a Btx-Alexa594 (bungarotoxin-Alexa594) conjugate (red). (B) Section of the single Z-stack slice showing a mixed population of myoblasts and myotubes. (C) Myoblasts (white outline) are identified as AChR positive cells containing 3 or more nuclei, while myoblasts (dotted white outline) contain 2 or less. Fusion indexes are calculated as the fraction of total nuclei contained in myotubes.
